# A phylogenetic method to perform genome-wide association studies in microbes that accounts for population structure and recombination

**DOI:** 10.1101/140798

**Authors:** Caitlin Collins, Xavier Didelot

## Abstract

Genome-Wide Association Studies (GWAS) in microbial organisms have the potential to vastly improve the way we understand, manage, and treat infectious diseases. Yet, GWAS methods established thus far remain insufficiently able to capitalise on the growing wealth of bacterial and viral genetic sequence data. Facing clonal population structure and homologous recombination, existing GWAS methods struggle to achieve both the precision necessary to reject spurious findings and the power required to detect associations in microbes. In this paper, we introduce a novel phylogenetic approach that has been tailor-made for microbial GWAS, which is applicable to organisms ranging from purely clonal to frequently recombining, and to both binary and continuous phenotypes. Our approach is robust to the confounding effects of both population structure and recombination, while maintaining high statistical power to detect associations. Thorough testing via application to simulated data provides strong support for the power and specificity of our approach and demonstrates the advantages offered over alternative cluster-based and dimension-reduction methods. Two applications to *Neisseria meningitidis* illustrate the versatility and potential of our method, confirming previously-identified penicillin resistance loci and resulting in the identification of both well-characterised and novel drivers of invasive disease. Our method is implemented as an open-source R package called treeWAS which is freely available at https://github.com/caitiecollins/treeWAS.

## Introduction

Owing to rapid progress in sequencing technologies, the accumulation of microbial genome sequences has begun to outpace the development of statistical and computational tools for their analysis. As a result, opportunities to reduce the global burden of infectious disease are missed, and infectious diseases remain accountable for 15% of worldwide annual mortality [1]. Moreover, as globalisation continues to increase the rate and scope of human interaction, with each other and with animals, this process will likely be accompanied by parallel change in the spread and evolution of infectious pathogens [2–5]. Discovering the genetic basis of microbial traits would offer key insights into the biological mechanisms underlying infectious diseases, and would improve our ability to develop drugs and vaccines, target treatments, build predictive tools, benefit from surveillance, and enhance public health.

Genome-wide association studies (GWAS) can be used to make these inferences, linking genotype to phenotype by testing for statistical associations between the two. GWAS have already become a tool of choice in human genetics, since the publication of the first such studies in the early 2000s [6–9], leading to the identification of over 11,000 trait-associated single nucleotide polymorphisms (SNPs) in humans [10]. It has been anticipated that by applying GWAS methods to microbes, similar discoveries could be made [11]. Indeed, although the advent of GWAS in microbes has been relatively recent, promising results can already be seen in the literature to date [12–25]. By contrast to GWAS in humans, however, microbial association mapping remains a technical challenge in search of an optimal methodological approach.

The purpose of GWAS, in microbes as in humans, is to identify statistically significant associations that may indicate the presence of a causal relationship between genotype and phenotype. Equally important is the inverse aim: to reject spurious associations arising from confounding factors. In microbes, smaller genome sizes and the ability to manipulate these genomes in the laboratory may improve the power and computational ease of GWAS and facilitate the confirmation of candidate loci [11]. On the other hand, microbial association studies must overcome a multiplicity of confounding factors, such as, the stronger population structure that results from clonal reproduction [26], widespread linkage disequilibrium interrupted unpredictably by homologous recombination [27], diversity in genetic content [28], and variability in the phenotypic probability distribution for a given genotype [29].

Most microbial GWAS analyses to date have made an effort to control for the confounding potential of population structure. The strength of this confounding effect increases both with the degree to which allele frequencies differ between subpopulations in a sample and the extent to which phenotypic states cluster within these lineages or clades [30,31]. Cluster-based methods [32] and dimension reduction techniques [33,34] have been adopted to account for population structure in microbial association studies [16,19,21,23–25,34,35]. Such methods, however, to the extent that they control for population structure also undermine the power to detect associations [24]. Furthermore, because the degree of phenotypic clustering is not factored into the analysis, they cannot appropriately evaluate the probability that population substructure will give rise to spurious associations. Methods that rearrange the phenotype to assess significance face the same limitation [16,22,25]. Pairwise methods account for fine scale genetic differences and phenotypic clustering, but at the cost of discarding large volumes of valuable data, thus reducing the power to detect associations [14,22]. Hence, despite the adoption of various strategies, the scope of relatedness in clonal organisms remains a particular challenge to microbial association studies.

Fortunately, clonality also enables the adoption of a phylogenetic solution [12,13]. Phylogenetic trees allow for the detailed identification of genetic relationships, not only at the level of population clusters, but also at the resolution of subpopulations and individual relationships. Adopting a phylogenetic approach does not require evolution to be treated as purely clonal, nor that recombination must be ignored, since the effect of recombination events can be considered within a phylogenetic framework [26,36]. Nor do they require any loss of information, provided pairwise techniques are not used. Phylogenetic approaches are by far the most popular method to describe microbial population structure, and therefore they are a natural option to control for population structure when performing GWAS in microbes.

Here we propose a new phylogenetic approach to GWAS called treeWAS that is able to overcome many of the limitations of existing microbial GWAS approaches. Within our analytical pipeline, data simulation based on parameters of the empirical dataset under analysis allows us to account for the composition of the genetic dataset and the population structure of the sample. In addition, by using the empirical homoplasy distribution to guide the simulation of null genetic data, we are able to control objectively for the confounding effects of recombination. We adopt multiple complementary scores of association that are able to handle both binary and continuous phenotypes. Applied in combination, these scores are designed to enhance statistical power and to improve detection of associations underlying subtle and complex phenotypes, such as host association or invasiveness. We present the results of rigorous testing on simulated datasets, and compare performance with alternative approaches, including cluster-based and dimension-reduction methods. We also demonstrate how treeWAS responds to varying levels of recombination and contrast this to previous methods. We show that treeWAS provides both specificity and power in a wide range of settings, and consistently offers the best overall performance. Finally, we present two applications to real data from *Neisseria meningitidis*. First, we investigate penicillin resistance and demonstrate that our approach can confirm known resistance loci. Second, we examine invasive disease, which reveals both previously characterised and novel invasiveness factors and illustrates the ability of our methodology to identify associations when applied to complex phenotypes.

## Materials and methods

### Overview of the treeWAS method

Our central aim is to delineate true signals of association from a noisy background of spurious associations. To accomplish this, our method uses the simulation of a null genetic dataset to establish whether high association score values in the empirical dataset under analysis are likely to be truly significant or may, in fact, arise by chance as a result of confounding factors found in the empirical dataset. We characterise the evolutionary parameters of the empirical dataset and use these to generate the simulated genetic dataset, which represents the null hypothesis of no association. This null dataset resembles the empirical dataset in both genetic composition and population structure, but does not have any true association with the phenotype. By comparing associations in the empirical and simulated datasets, we are able to determine which signals of association have sufficient statistical and evolutionary support. This approach makes use of all information contained in the dataset, as well as that inferred in phylogenetic and ancestral state reconstructions. We aim to maintain strict control over the number of false positive findings. This makes possible the application of multiple complementary tests of association, which increases the power to detect associations.

### Implementation

The treeWAS approach is implemented in the following steps:

1. **Phylogenetic reconstruction** can be performed within treeWAS by distance-based [37–40] or maximum-likelihood (ML) [41] methods. However, where recombination is expected to distort the clonal genealogy, it is recommended that users provide a tree previously reconstructed by a recombination-aware approach [36,42,43]. Tools are provided for integration with ClonalFrameML [42].
2. Computation of the homoplasy distribution, containing site-specific numbers of substitutions drawn from the empirical dataset, is performed with the Fitch parsimony algorithm [44].
3. **Simulation of null genetic data** enables the delineation of true associations from spurious associations. We compare the relationships between genotype and phenotype at all loci in the real data to those in a simulated dataset that embodies only potentially confounding factors. Simulation of this “null” genetic dataset is guided by three parameters: (i) the phylogenetic tree, (ii) the homoplasy distribution, (iii) the number of loci to be simulated, *N_sim_*, which is recommended to be at least ten times the number of biallelic sites in the empirical dataset. Each of the *N_sim_* loci is simulated along the phylogenetic tree, from root to tips, undergoing a number of substitutions drawn from the homoplasy distribution on branches selected randomly with probabilities proportional to branch length. The original phenotype is maintained across the leaves. By retaining the phylogenetic tree and the distribution of phenotypic states, but reassigning substitutions to new branches, we are able to produce a simulated dataset that resembles the empirical dataset in population structure and genetic composition, including the effects of mutation and recombination. Associations due to confounding factors, but no true associations with the phenotype, are thus recreated by the simulation process.
4. **Ancestral character estimation** is required prior to association testing. A marginal reconstruction of the ancestral states of both genotype and phenotype must be performed via parsimony [45] (or ML [46,47] for continuous phenotypes).
5. **Association testing** is performed by applying three independent tests of association to all loci (see below). Associations between simulated loci and the empirical phenotype are first measured, allowing for the identification of a null distribution of association score statistics under the null hypothesis of no association. Associations between empirical loci and the phenotype are then measured and evaluated with reference to this null distribution.
6. **Identification of significance threshold and associations** proceeds by drawing a threshold in the upper tail of the null distribution, at the value corresponding to a base p-value (e.g., *p* = 0.01) that has been corrected for multiple testing (via Bonferroni correction by default, though False Discovery Rate is also implemented). Among the set of empirical association scores, all values that exceed this threshold are deemed to be statistically significant associations and, thus, candidates for true biological association, pending subsequent confirmatory analyses.

### Tests of association

The design of treeWAS, particularly its use of the null distribution, enables strict control over the false positive rate. This presents the opportunity for power to be augmented by applying multiple independent tests of association. We therefore measure the association between each genetic locus and the phenotype with three separate tests, described below and illustrated in Fig 1. All three tests are applicable to any form of binary genetic data, including SNPs, indels, and gene presence or absence matrices, and can be used on both binary and continuous phenotypic data. An association deemed significant by any one test is a suitable candidate for further investigation. Significance according to a second or third test is not required, but may provide further support for a finding. The three scores have been designed to complement each other and to recognise distinct, if overlapping, patterns of association. The following notations are used to describe the three association scores. 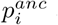 and 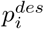 denote the phenotypic state at ancestral and descendant node of branch *i*, respectively. 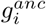 and 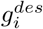 denote respectively the genotypic state at the ancestral and descendant node of branch *i. n* and *n_b_* denote respectively the number of leaves and branches on the phylogenetic tree.

**Fig 1.**
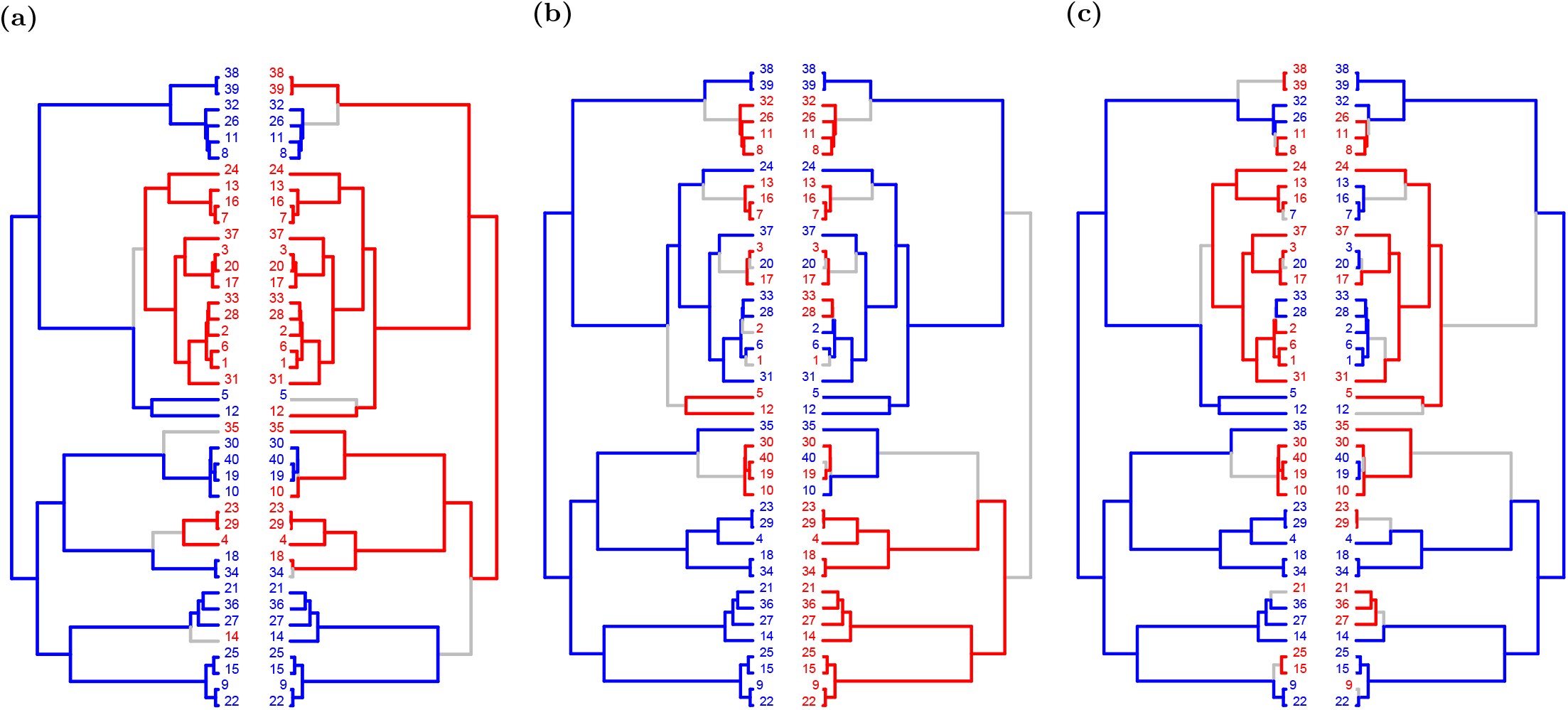
Evolutionary scenarios detected by treeWAS scores. The three complementary tests of association in treeWAS assign high scores to different patterns of association, examples of which are illustrated above. Each panel displays the phenotype (left) and the genotype of one associated locus (right), with binary states plotted along the tips of the phylogenetic tree (N = 40) and reconstructed ancestral states indicated along the branches of the tree (blue = 0, red = 1, grey = substitution). **A**: Score 1 considers only the tips of the tree and, with uniform association in all but six terminal nodes in this scenario, it would receive a Score 1 value of 0.7 by Eq 1. Conversely, Scores 2 and 3 would obtain low values because no simultaneous substitutions or maintained patterns of association are observed among the reconstructed ancestral states. **B**: Score 2 counts the number of branches containing a substitution in both genotype and phenotype, resulting in a Score 2 value of 5 for this scenario. Score 2 is not penalised for the lack of simultaneous substitutions in the lower major clade, whereas Scores 1 and 3 would receive low values for being out of association in half of this tree. **C**: Score 3 detects subtler patterns of association and uses the reconstructed ancestral states to infer the maintenance of association across the branches of the phylogenetic tree, resulting in a Score 3 value of 10 to this scenario. Meanwhile, the absence of terminal association and lack of simultaneous substitutions mean that this pattern receives values of zero by both Score 1 and Score 2.

**Score 1**, the “Terminal Score”, measures sample-wide association across the leaves of the phylogenetic tree. For a binary phenotype, this score is equivalent to counting the four terminal state combinations, with and without the phenotype, and with and without the genotype, as previously proposed [48]. Generalizing to continuous phenotypes gives:

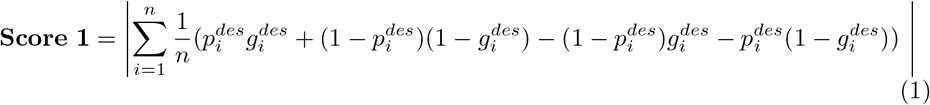

This score is blind to the number of origins of association, but also robust to the potential downstream effects of incorrectly estimating the phylogenetic tree or ancestral states of the phenotype or genotype, as illustrated by Fig 1A.

**Score 2**, the “Simultaneous Score”, measures the degree of parallel change in the phenotype and genotype across branches of the tree. For a binary phenotype with a parsimonious ancestral state reconstruction, as in Fig 1B, this means counting the number of branches containing a simultaneous substitution in genotype and phenotype [13]. A more general definition is given by:

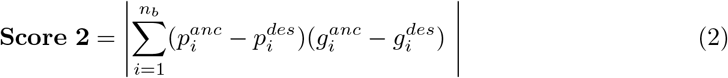

Unlike Score 1, Score 2 is able to use information contained in the tree structure and set of ancestral character states towards the detection of significant associations. Moreover, because Eq 2 imparts a cumulative character (simultaneous substitutions increase the score, but branches where one or no variable changes do not decrease it), significance by Score 2 does not require sample-wide association. Score 2 may therefore be able to detect loci giving rise to the phenotype through complementary pathways, in addition to identifying loci whose associations with the phenotype persist across the tree.

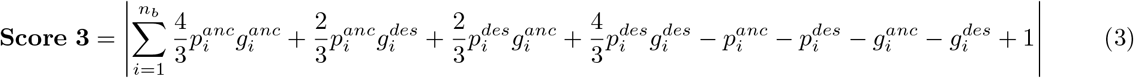

**Score 3**, the “Subsequent Score”, measures the proportion of the tree in which the genotype and phenotype co-exist. It is the mathematical solution to the integral of an association score along all points of the phylogenetic tree (see S1 Appendix):

Score 3 considers the maintenance of allelic enrichment in a given phenotypic state, as well as the change in both genotype and phenotype. Should association arise by a substitution in one variable being followed on a subsequent branch by a substitution in the other, as in Fig 1C, Score 3 will incur no penalty for the lack of simultaneous change and will capture the downstream association in so far as it is maintained. This score should therefore be sensitive to subtler and more probabilistic patterns of association. For example, Score 3 may be effective in the case of host association, where genetic adaptation contributes both before and after host switching, for example, by increasing affinity for a new host or by offering compensatory fitness advantages in a new environment [48].

### Assessment of performance on simulated data

To evaluate the performance of treeWAS, we applied it and six alternative methods to 240 simulated datasets. Three separate approaches were used to simulate 80 datasets each. Approaches differed only in the nature of the simulated associations between genotype and phenotype. We present one approach below and the other two in S2 Appendix and S3 Appendix.

Each simulated dataset contained 100 individuals and 10,000 binary loci, of which ten loci were associated with a binary phenotype. The non-associated loci were simulated using homoplasy distributions corresponding to four recombination rates (0, 0.01, 0.05, 0.1) (see S4 Appendix and S1 Fig). Genetic data was simulated along randomly generated coalescent trees. For each simulated non-associated locus, the number of substitutions was drawn from the homoplasy distribution and assigned to branches of the phylogenetic tree with probabilities proportional to branch lengths. The ten associated loci were generated together with the phenotype according to an instantaneous transition rate matrix, *Q*, which controls the rates of transition between all four possible combinations of a binary genetic locus, *G*, and the binary phenotype, *P*, (i.e., *G*_0_*P*_0_, *G*_0_*P*_1_, *G*_1_*P*_0_, *G*_1_*P*_1_) between an ancestral node (in the rows) and a descendant node (in the columns):

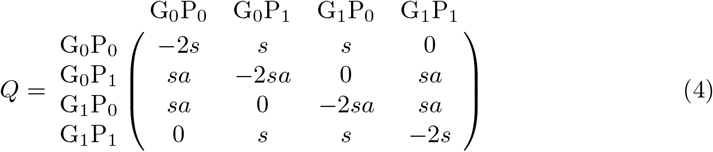

The *Q* matrix is parameterised by *s*, which controls the baseline substitution rate and applies to all columns, and *a*, an association factor that establishes the preference for one form of association (*G*_0_*P*_0_, *G*_1_*P*_1_) over the opposite (*G*_0_*P*_1_, *G*_1_*P*_0_). The parameter *s* is divided by the sum of the branch lengths before building *Q*. In all simulations, initial parameters were set to *s* = 20 and *a* = 10. To identify the probabilities of transition for a branch of length *l*, the instantaneous transition rate matrix, *Q*, is converted into a matrix of probabilities, *P* = exp(*Ql*), via matrix exponentiation, which takes into account the length *l* of the branch in question.

### Comparison with other GWAS methods

To each of the simulated datasets, in addition to treeWAS, we applied six alternative GWAS methods. The Fisher’s exact test, and the χ^2^ test available in PLINK version 1.07 [49] were used as benchmarks to demonstrate what results would be found by two standard tests of association without population structure control. The PLINK χ^2^ test with Genomic Control (GC), has been used in bacterial GWAS [21] and provided a simple solution to population structure. Principal Components Analysis (PCA) and Discriminant Analysis of Principal Components (DAPC) represent more advanced and popular approaches to correcting for population structure [33,34]. PCA is the “gold standard” method used in human GWAS [30, 50] and DAPC has been proposed as a potential improvement on PCA [34]. Both have been used in microbial GWAS [19,23–25,34]. We followed the standard protocol used in human genetics and corrected for ancestry by regressing along the significant Principal Components (PCs) of PCA or DAPC (see S5 Appendix), identifying significant associations via χ^2^ test [30]. The Cochran-Mantel-Haenszel (CMH, [51]) provided an alternative, cluster-based approach. The CMH test works directly with *K* population clusters by adopting a stratified 2x2x*K* design and has been used in bacterial GWAS [16,21,35]. Results of methods applied to the simulations described in S2 Appendix are presented in S2-S7 Figs. Results for the simulations described in S3 Appendix are presented in S8-S13 Figs. Results for the simulations described above are presented below and in S14-S17 Figs.

## Results and Discussion

### Assessment of treeWAS performance on simulated data

The performance on simulated datasets was evaluated along four metrics: the False Positive Rate (FPR), Sensitivity, Positive Predictive Value (PPV), the proportion of results that are true positives, and the F1 Score, which is the harmonic mean of Sensitivity and PPV [52]. Our approach performed well along all four metrics. Fig 2A shows that treeWAS was able to consistently achieve a FPR of zero, indicating tight control over multiple confounding factors. We see from Fig 2B that this conservative approach produces moderate sensitivities in the three individual association scores within treeWAS, but that the contribution of true positive findings by each score gives treeWAS a very high sensitivity overall. Notably, although Score 2 generally achieved higher sensitivity than Scores 1 and 3, it was not always the leading contributor to the cumulative sensitivity of treeWAS in the analysis of these simulated datasets. This highlights the value of using multiple complementary measures to identify associations. Importantly, cumulative benefits in sensitivity were not undermined by cumulative reductions in PPV. Fig 2C reflects the fact that in most cases the total number of false positives found by treeWAS was zero or one. Overall, the high performance of our approach, as indicated by the composite F1 score in Fig 2D, provides strong support for the strategy adopted by treeWAS.

**Fig 2.**
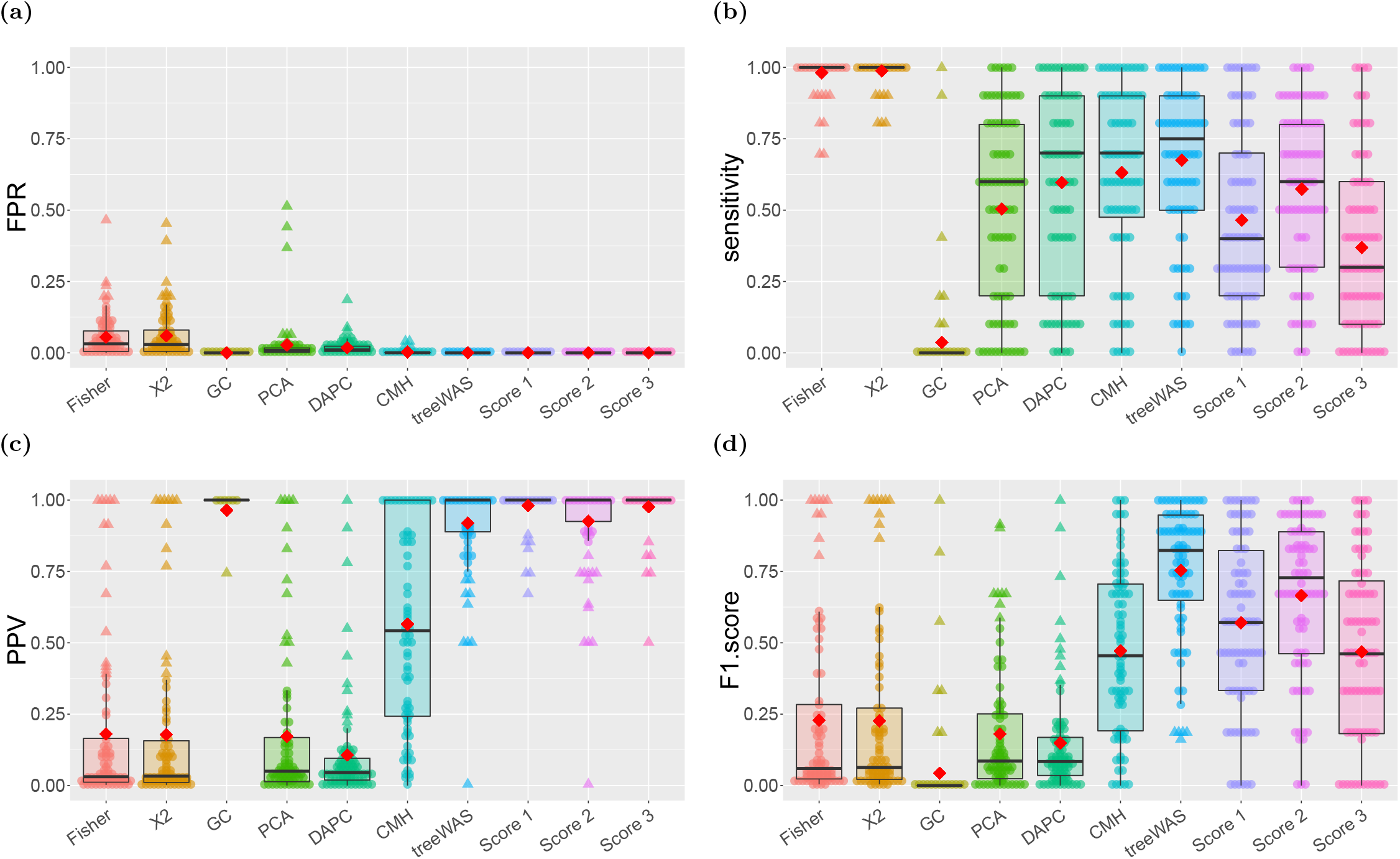
Performance by association test. The performance on simulated datasets for the six comparator GWAS methods and treeWAS, alongside its three association tests individually, is summarised along the four metrics of evaluation. Box plots display the median and interquartile range, red diamonds indicate the mean, and individual dots represent results for one of the 80 simulated datasets. **A**: False Positive Rate. **B**: Sensitivity. **C**: Positive Predictive Value. **D**: F1 Score.

In addition to providing a thorough control over population structure, treeWAS was able to account explicitly for the varying confounding effect of recombination. In the simulated datasets analysed above, the phenotype and associated loci underwent between four and 23 binary state substitutions as a result of the probabilistic simulation process, with an average of 14 substitutions per tree. As such, the probability of chance correlation with the resulting pattern of phenotypic clustering increased as the number of substitutions among non-associated loci was elevated to similar levels by recombination (see S1 Fig). Fig 3A shows that the FPR of treeWAS remained consistently low as the recombination rate was increased. Because data simulation within treeWAS is guided by the empirical homoplasy distribution, the elevated risk of chance association due to recombination was accounted for in the null distribution. Fig 3B illustrates a second implication of this feature: in this analysis, sensitivity decreased with increasing recombination, as treeWAS could no longer attribute significance to some more weakly associated loci when similar patterns of association were likely to occur by chance. Taking into account the parameters of the data causes the impact of recombination on the sensitivity of treeWAS to vary by context (see S3 Fig and S9 Fig). This data-dependent behaviour is necessary to keep FPR at a minimum (Fig 3A). We do nevertheless see in Fig 3C a slight decline in the PPV of treeWAS, indicating a shift from an average of zero to one false positive findings with increasing recombination. Ultimately, as the F1 score in Fig 3D demonstrates, the approach adopted by treeWAS not only produces good overall performance, but by accounting for recombination, it is able to maintain good performance across a range of backgrounds, from purely clonal to frequently recombining.

**Fig 3.**
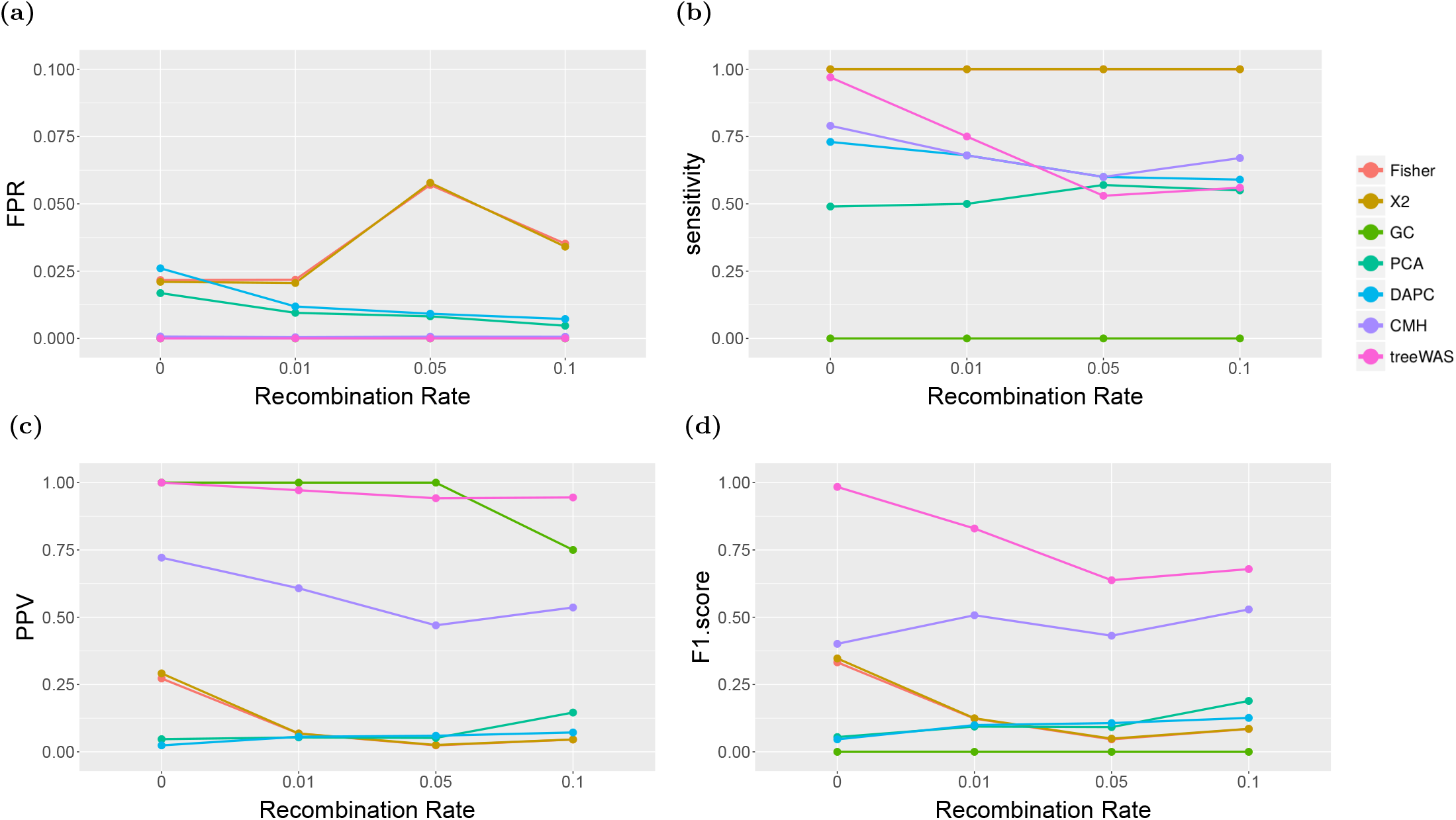
Performance by recombination rate. Interquartile mean performance by GWAS method and recombination rate is plotted along four statistics. **A**: False Positive Rate. **B**: Sensitivity. **C**: Positive Predictive Value. **D**: F1 Score.

### Comparison with other GWAS methods

Having described the performance of treeWAS on simulated data, we now compare it with the performance of alternative methods. Fig 2A reveals that only treeWAS and the conservative GC approach consistently rejected all false positive findings. PCA, DAPC, and the CMH test reduced FPR below the level incurred with no correction for population structure, but still returned undesirably high volumes of false positives. Fig 2A thus suggests that even the most popular dimension-reduction and cluster-based methods do not sufficiently correct for population structure in microbial contexts.

All of the non-phylogenetic approaches to correcting for population structure inherently simplify the extensive genetic relationships between isolates. The k-means clusters used in the CMH test, and the principal components of PCA and DAPC have been previously shown to correspond to the major clades and genealogical divisions of a phylogenetic tree [19,23,53]. In PCA, DAPC, and the CMH test, therefore, the user must make a conceptual delineation, at a given height on this tree, between what will and will not be considered parts of the population structure. Our approach, by contrast, works directly with the whole phylogenetic tree, retaining the information it provides at all levels of the clonal population structure. In addition, because the distribution of the phenotype is observed along the tips of the phylogeny but less clear in clusters and PCs, treeWAS is able to account directly for the degree of correlation observed between population structuring alleles and phenotypic clusters. The design of treeWAS therefore allows it to determine what degree of association is unlikely to have arisen by chance, given the evolutionary history inferred. For these reasons, our phylogenetic approach provides a more natural and complete solution to the problem posed by population structure, which drives the FPR gap in Fig 2A between treeWAS and its competitors.

Building on the foundation of low FPR, treeWAS is able to enhance power by drawing on the cumulative findings of multiple tests of association. The non-phylogenetic alternatives, by contrast, face an inherent trade-off between sensitivity and specificity. Fig 2B and C show that, consequently, none of these approaches offered simultaneously a high PPV and high sensitivity. The only non-phylogenetic method with an acceptably high PPV was GC (Fig 2C), but it also had the lowest sensitivity (Fig 2B) meaning that in our simulations it almost never found any association, correct or incorrect. PCA and DAPC, despite being too permissive of false positives, achieved only moderate sensitivities, considerably lower than treeWAS (Fig 2B). These methods undermine sensitivity by discounting higher-order lineage effects and proceed with the assumption that remaining genetic variation is ancestrally homogenous [23,54]. Because these methods reduce power by eliminating variation with every additional PC, increasing the number of selected PCs in an effort to lower FPR in this study resulted in a complete loss of sensitivity. Our results suggest that when PCA and DAPC are used in microbial GWAS, depending on the population structure and the effect size of associations, a satisfactory trade-off between sensitivity and specificity may be unattainable.

The CMH test more effectively managed the sensitivity-specificity trade-off by applying a more conservative stratified test of association to the genetic data matrix without regressing out any relevant information. In fact, we see from Fig 2B that the sensitivity of the CMH test was only slightly lower than that of treeWAS. The CMH test also had better PPV than PCA and DAPC (Fig 2C), although it fell well below that of treeWAS. Indeed, with an mean PPV of 0.56, almost half of the results identified by the CMH test were false positives, whereas the PPV of treeWAS indicates that 92% of our findings were correct. Overall, the F1 score (Fig 2D) of CMH was similar to that of the lowest-performing individual association test within treeWAS. Yet, the CMH test achieves high F1 score values by adopting the less stringent approach of favouring high sensitivity over high PPV. In practice, it may be preferable to incur modest sensitivity losses so that the number of false positive findings may be kept at a minimum.

Additionally, Fig 3B and C show that because the CMH test is naive to recombination, its behaviour differed markedly from that of treeWAS. In response to increasing recombination, the CMH test maintained relatively stable sensitivity, but experienced a decrease in PPV as the number of false positive findings increased, demonstrating a lack of control for recombination. Furthermore, although the composite F1 scores in Fig 3D appear to narrow with increasing recombination, it is important to note the practical implications of the trade-offs being made by treeWAS and the CMH test. Even at the highest recombination rate examined, the CMH test identified only one more true positive than treeWAS. On the other hand, treeWAS found less than one false positive on average, while the CMH test found as many false positives as true positives (Fig 3C). Increasing recombination in other simulations caused the F1 score gap between the CMH test and treeWAS to increase or remain unchanged (S3 and S9 Fig).

Overall, the comparison of methods in Fig 2 and 3 indicates that the performance of the non-phylogenetic methods was limited by multiple factors: the focus on higher level population structure, the inability to control for its confounding effects in sufficient detail, the necessary trade-off between sensitivity and PPV, and the poor response to varying rates of recombination. By deliberately avoiding all of these pitfalls, the design of treeWAS achieved stronger performance on these simulated datasets.

### Application to *Neisseria meningitidis*: identifying penicillin resistance factors

To determine whether our approach could confirm previously-identified associations, and illustrate its applicability to both binary and continuous phenotypes, we applied treeWAS to a dataset of *N. meningitidis* isolates with a penicillin resistance phenotype. We used the *Neisseria* Bacterial Isolate Genome Sequence Database (BIGSdb accessible at https://pubmlst.org/neisseria/, [55]) to download 171 *N. meningitidis* sequences from serogroup B (see S1 File), extracting both the core SNPs (166,848 SNPs) and an accessory gene presence or absence matrix (2,808 genes). We reconstructed the phylogenetic tree from whole-genome sequences with ClonalFrameML to account for recombination [42]. *N. meningitidis* has a high recombination rate, though recombination in *N. meningitidis* is not so rampant as to entirely obscure the clonal genealogy, as would be the case for example in *Helicobacter pylori* [56]. Because treeWAS accounts explicitly for the confounding effects of recombination, our approach was appropriate for this context, provided recombination-aware phylogenetic methods were used [57].

The penicillin resistance phenotype was analysed in two ways: as a binary and as a continuous variable. The binary phenotype was categorised according to the penicillin minimum inhibitory concentration (MIC), defining susceptible as MIC ≤ 0.06 and resistant as MIC > 0.06. The continuous phenotype was defined as the ranks of the MIC values, rather than the MIC values themselves, whose distribution was highly skewed and uninformative.

Analysis of the accessory gene presence or absence data did not result in the identification of any gene significantly associated with either the binary or continuous penicillin resistance phenotype. However, application of treeWAS to the set of core SNPs led to the identification of many significant loci. Analysis of the binary penicillin resistance phenotype resulted in the identification of 162 significant SNPs, all of which were located in the well-characterised NEIS1753 (*penA*) gene, encoding penicillin-binding protein 2 (see S2 File and S18 Fig). This finding is consistent with the literature, which indicates that penicillin resistance in *N. meningitidis* occurs when altered forms of a penicillin-binding protein (PBP) are produced [58]. Previous work also indicates that the resistance phenotype, and the mosaic structure of *penA*, arise via homologous recombination [59,60]. It is therefore natural that our analysis uncovered significant associations among SNPs in this gene rather than in other core or accessory genes. Indeed, the alignment displayed in S18A Fig is consistent with previous accounts in the literature describing uniformity in *penA* sequences among susceptible isolates [61] and considerable diversity among those in resistant isolates [62].

Analysis of the continuous penicillin MIC phenotype returned 30 significant SNPs (see S3 File and S19 Fig). The majority of these were also located in the *penA* gene, although SNPs were also identified in three additional genes. In the presence of antibiotics, many loci not essential to the resistance phenotype may confer a slight selective advantage [63]. For example the UDP-N-acetylmuramoylalanyl-D-glutamate–2, 6-diaminopimelate ligase is involved in cell wall formation via peptidoglycan synthesis, the process targeted by penicillin [60]. The two additional genes in which significant SNPs were identified have roles in stress response and DNA damage repair, which may not be directly related to penicillin resistance but which may instead confer a minor fitness advantage that would slightly increase MIC values. It should be noted that although the classification scheme we adopted designated MIC > 0.06 as “resistant”, these isolates would in fact usually be classified as of “intermediate resistance”, as they did not exceed the standard resistance threshold of MIC > 1 (see S1 File). In light of the narrow range of MIC values in the sample, it is remarkable that treeWAS was nonetheless able to identify significant associations. By analysing resistance as a continuous variable, our approach was able to retain all of the phenotypic information available. Hence, in spite of the relatively small phenotypic effect observed, treeWAS not only retained the power to detect the central PBP gene, *penA*, but it gained sensitivity to significant SNPs in three additional genes.

### Application to *Neisseria meningitidis*: identifying drivers of invasive disease

The overall design of treeWAS, in particular the implementation of the three association scores in Eqs 1–3, was developed with the aim of detecting genetic loci associated with subtle and complex phenotypes which may not be entirely determined by genetic factors [29]. To illustrate this, we applied treeWAS to a separate *N. meningitidis* dataset, with the more challenging phenotype of invasive disease versus carriage. Invasiveness is determined more probabilistically than penicillin resistance, on the basis of both pathogen genetics and external factors, such as host immunity [64].

From the *Neisseria* BIGSdb database, we downloaded 129 European *N. meningitidis* sequences from serogroup C (see S4 File), including both core SNPs (115,386 SNPs) and accessory gene presence or absence data (2,809 genes). ClonalFrameML was used to reconstruct the phylogenetic tree from whole-genome sequences while accounting for recombination [42].

In the analysis of the accessory gene presence or absence data, treeWAS identified 12 genes associated with carriage or invasiveness (Fig 4, Table 1). Three genes were found to be associated with invasive disease, and the role of each was confirmed by the literature. *NadA* (Neisserial adhesin A) has well-characterised roles in virulence, enabling adhesion, colonisation, and invasion of mucosal cells [65,66]. *MafA2*, another adhesin, plays a similar role in pathogenic *Neisseria* [67,68]. Epidemiological evidence and rat models have also linked the haemoglobin receptor protein, *hmbR*, to invasive disease in *N. meningitidis* [69,70]. Moreover, as this gene is highly conserved, *hmbR* may be a good target for vaccine development [71].

**Fig 4.**
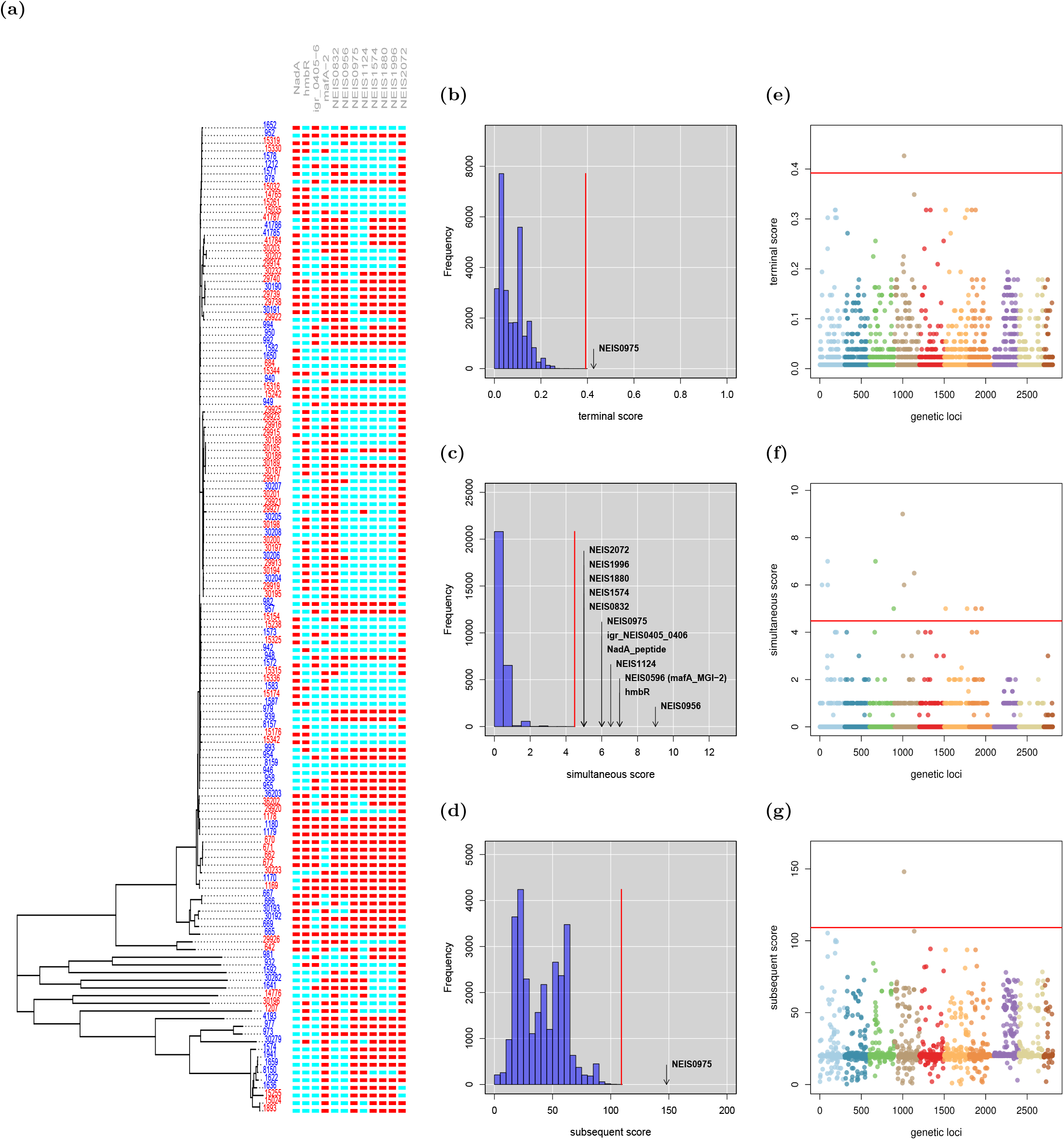
Invasive disease in the *N. meningitidis* accessory genome. treeWAS identified 12 genes associated with invasive disease. **A:** At left, the clonal genealogy reconstructed with ClonalFrameML, and terminal phenotype (blue = carrier, red = invasive). At right, an alignment of the 12 significant genes (blue = gene absence, red = gene presence). **B-D**: Null distributions of simulated association scores for (B) Score 1, (C) Score 2, (D) Score 3, a significance threshold (red), above which real associated genes are indicated. **E-G**: Manhattan plots for (E) Score 1, (F) Score 2, (G) Score 3 showing association score values for all genes, a significance threshold (red), above which points indicate significant associations.

**Table 1.** Genes associated with invasive disease in *N. meningitidis*.

| Gene | Gene product |
| --- | --- |
| NadA_peptide | *NadA* peptide |
| hmbR | Haemoglobin receptor protein |
| igr_NEIS0405_0406 | intergenic region between NEIS0405 and NEIS0406 |
| NEIS0596 (*mafA*_MGI-2) | *MafA-2* adhesin |
| NEIS0832 | hypothetical protein |
| NEIS0956 | cell-surface protein |
| NEIS0975 | hypothetical protein |
| NEIS1124 | hypothetical protein |
| NEIS1574 | DNA transport competence protein |
| NEIS1880 | DNA transport competence protein |
| NEIS1996 | DNA transport competence protein |
| NEIS2072 | putative periplasmic protein |

These genes were identified as significantly associated with invasive disease when treeWAS was applied to 129 accessory genome gene presence-or-absence sequences from *N. meningitidis* serogroup C.

We also identified nine genes whose presence was associated with Neisserial carriage. These included the cell-surface protein encoded by NEIS0956 and the DNA transport competence proteins encoded by NEIS1574, NEIS1880, and NEIS1996, which enable genetic transformation [72,73]. These genes may confer an adaptive advantage to *N. meningitidis* that enables immune evasion via surface modulation, and favours colonisation and survival in the nasopharyngeal niche [74]. This relationship is not entirely clear, however, as non-pathogenic carriage remains incompletely characterised at a molecular level, despite being a fundamental element of the Neisserial life cycle [75].

In the analysis of core SNPs, treeWAS identified seven associated loci (Fig 5, Table 2). Among these, the *porA* gene is well known for encoding a surface protein that drives hyperinvasivity in *N. meningitidis* [76–79]. Likewise, *gapA-2* may facilitate the adhesion to and invasion of host tissues [80]. As the genetic basis of invasiveness in *N. meningitidis* is not yet fully understood, we anticipate that future work will elucidate the roles of other loci in Table 2.

**Fig 5.**
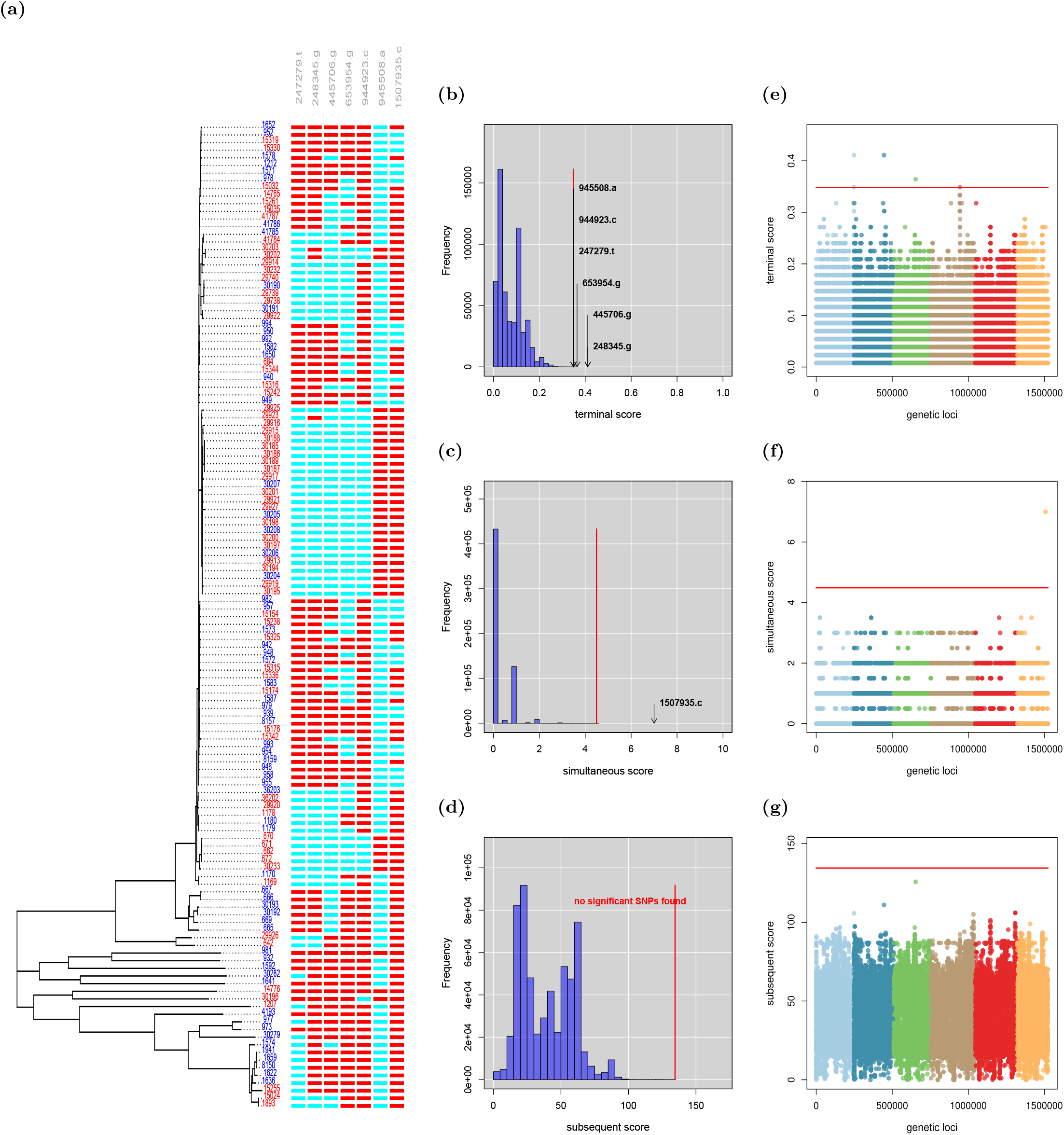
Invasive disease in *N. meningitidis* core SNPs. treeWAS identified 7 SNPs associated with invasive disease. **A**: At left, the clonal genealogy reconstructed with ClonalFrameML, and terminal phenotype (blue = carrier, red = invasive). At right, an alignment of the 7 significant SNPs (blue = allele 0; red = allele 1). **B-D**: Null distributions of simulated association scores for (B) Score 1, (C) Score 2, (D) Score 3, a significance threshold (red), above which real associated SNPs are indicated. **E-G**: Manhattan plots for (E) Score 1, (F) Score 2, (G) Score 3 showing association score values for all SNPs, a significance threshold (red), above which points indicate significant associations.

**Table 2.**
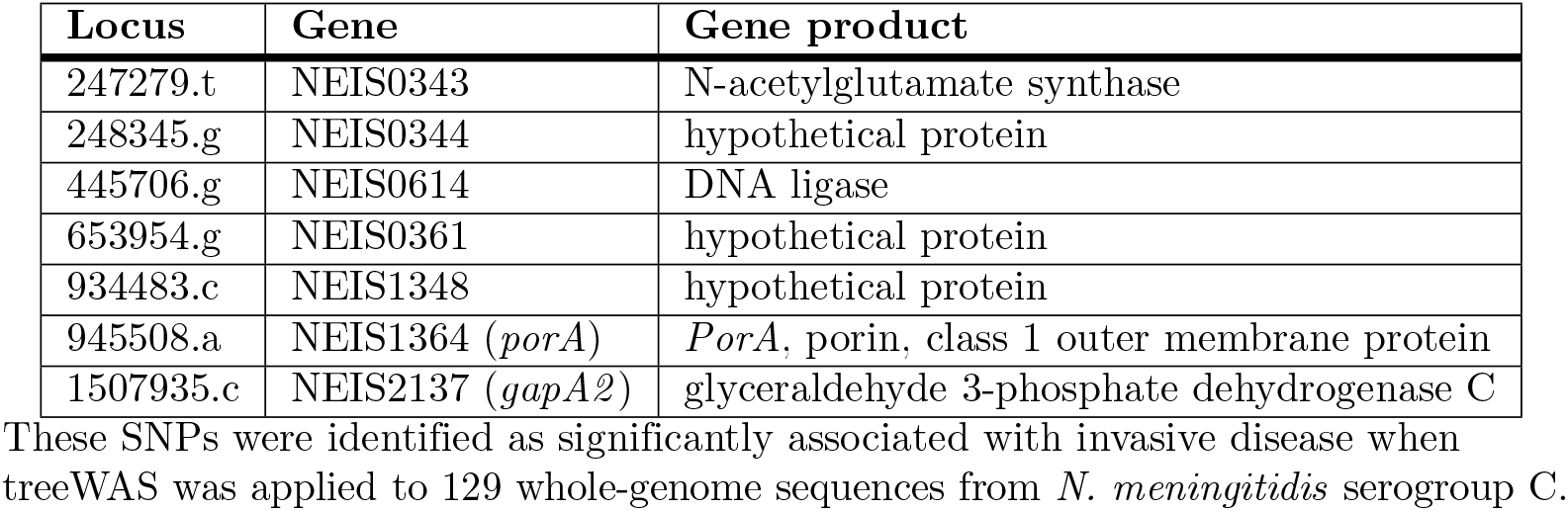
SNPs associated with invasive disease in *N. meningitidis*.

Overall, treeWAS was able to identify both previously-known and putatively novel genes and SNPs in significant association with the commensal or invasive phenotype in *N. meningitidis*. Subsequent analyses in the laboratory would be required to confirm that a true biological or causal relationship accompanies this statistical significance.

## Conclusions

Microbial GWAS has the potential to reveal many important features of microbial genomes. Application has however been so far hampered by a lack of well founded and thoroughly tested methodology. Here we proposed a new phylogenetic approach to microbial GWAS that is able to control for the disruptive effects of both population structure and recombination, whilst still retaining a high statistical power to detect real associations. Application to both simulated and real datasets demonstrated that our method is accurate, efficient and versatile, being able to detect associations in both the core and pan-genome and for both categorical and continuous phenotypic measurements. We have implemented our approach in a user-friendly R package, treeWAS, which is freely available for public use at https://github.com/caitiecollins/treeWAS.

## Acknowledgments

This work was support by BBSRC grant BB/L023458/1 and NIHR grant HPRU-2012-10080. We would like to thank David Aanensen, Christophe Fraser and Daniel Wilson for useful discussions.

## References

1. WHO, “World health statistics. global health indicators: Cause-specific mortality and morbidity,” World Health Organisation, p. 72, 2015.

2. B. V. Lowder, C. M. Guinane, N. L. Ben Zakour, L. A. Weinert, A. Conway-Morris, R. A. Cartwright, A. J. Simpson, A. Rambaut, U. Nübel, and J. R. Fitzgerald, “Recent human-to-poultry host jump, adaptation, and pandemic spread of staphylococcus aureus,” Proc. Natl. Acad. Sci. U. S. A., vol. 106, pp. 19545–19550, 17 Nov. 2009.

3. C. M. Guinane, N. L. Ben Zakour, M. A. Tormo-Mas, L. A. Weinert, B. V. Lowder, R. A. Cartwright, D. S. Smyth, C. J. Smyth, J. A. Lindsay, K. A. Gould, A. Witney, J. Hinds, J. P. Bollback, A. Rambaut, J. R. Penadés, and J. R. Fitzgerald, “Evolutionary genomics of staphylococcus aureus reveals insights into the origin and molecular basis of ruminant host adaptation,” Genome Biol. Evol., vol. 2, pp. 454–466, 12 July 2010.

4. F. L. Kiechle, X. Zhang, and C. A. Holland-Staley, “The -omics era and its impact,” Arch. Pathol. Lab. Med., vol. 128, pp. 1337–1345, Dec. 2004.

5. M. T. G. Holden, L.-Y. Hsu, K. Kurt, L. A. Weinert, A. E. Mather, S. R. Harris, B. Strommenger, F. Layer, W. Witte, H. de Lencastre, R. Skov, H. Westh, H. Zemlicková, G. Coombs, A. M. Kearns, R. L. R. Hill, J. Edgeworth, I. Gould, V. Gant, J. Cooke, G. F. Edwards, P. R. McAdam, K. E. Templeton, A. McCann, Z. Zhou, S. Castillo-Ramírez, E. J. Feil, L. O. Hudson, M. C. Enright, F. Balloux, D. M. Aanensen, B. G. Spratt, J. R. Fitzgerald, J. Parkhill, M. Achtman, S. D. Bentley, and U. Nübel, “A genomic portrait of the emergence, evolution, and global spread of a methicillin-resistant staphylococcus aureus pandemic,” Genome Res., vol. 23, pp. 653–664, Apr. 2013.

6. J. Marchini, L. R. Cardon, M. S. Phillips, and P. Donnelly, “The effects of human population structure on large genetic association studies,” Nat. Genet., vol. 36, pp. 512–517, May 2004.

7. L. A. Weiss, J. Veenstra-Vanderweele, D. L. Newman, S.-J. Kim, H. Dytch, M. S. McPeek, S. Cheng, C. Ober, E. H. Cook, Jr, and M. Abney, “Genome-wide association study identifies ITGB3 as a QTL for whole blood serotonin,” Eur. J. Hum. Genet., vol. 12, pp. 949–954, Nov. 2004.

8. J. L. Haines, M. A. Hauser, S. Schmidt, W. K. Scott, L. M. Olson, P. Gallins, K. L. Spencer, S. Y. Kwan, M. Noureddine, J. R. Gilbert, N. Schnetz-Boutaud, A. Agarwal, E. A. Postel, and M. A. Pericak-Vance, “Complement factor H variant increases the risk of age-related macular degeneration,” Science, vol. 308, pp. 419–421, 15 Apr. 2005.

9. R. J. Klein, C. Zeiss, E. Y. Chew, J.-Y. Tsai, R. S. Sackler, C. Haynes, A. K. Henning, J. P. SanGiovanni, S. M. Mane, S. T. Mayne, M. B. Bracken, F. L. Ferris, J. Ott, C. Barnstable, and J. Hoh, “Complement factor H polymorphism in age-related macular degeneration,” Science, vol. 308, pp. 385–389, 15 Apr. 2005.

10. D. Welter, J. MacArthur, J. Morales, T. Burdett, P. Hall, H. Junkins, A. Klemm, P. Flicek, T. Manolio, L. Hindorff, and H. Parkinson, “The NHGRI GWAS catalog, a curated resource of SNP-trait associations,” Nucleic Acids Res., vol. 42, pp. D1001–6, Jan. 2014.

11. D. Falush and R. Bowden, “Genome-wide association mapping in bacteria?,” Trends Microbiol., vol. 14, pp. 353–355, Aug. 2006.

12. S. K. Sheppard, X. Didelot, K. A. Jolley, A. E. Darling, B. Pascoe, G. Meric, D. J. Kelly, A. Cody, F. M. Colles, N. J. C. Strachan, I. D. Ogden, K. Forbes, N. P. French, P. Carter, W. G. Miller, N. D. McCarthy, R. Owen, E. Litrup, M. Egholm, J. P. Affourtit, S. D. Bentley, J. Parkhill, M. C. J. Maiden, and D. Falush, “Progressive genome-wide introgression in agricultural campylobacter coli,” Mol. Ecol., vol. 22, pp. 1051–1064, Feb. 2013.

13. M. R. Farhat, B. J. Shapiro, K. J. Kieser, R. Sultana, K. R. Jacobson, T. C. Victor, R. M. Warren, E. M. Streicher, A. Calver, A. Sloutsky, D. Kaur, J. E. Posey, B. Plikaytis, M. R. Oggioni, J. L. Gardy, J. C. Johnston, M. Rodrigues, P. K. C. Tang, M. Kato-Maeda, M. L. Borowsky, B. Muddukrishna, B. N. Kreiswirth, N. Kurepina, J. Galagan, S. Gagneux, B. Birren, E. J. Rubin, E. S. Lander, P. C. Sabeti, and M. Murray, “Genomic analysis identifies targets of convergent positive selection in drug-resistant mycobacterium tuberculosis,” Nat. Genet., vol. 45, pp. 1183–1189, Oct. 2013.

14. M. Farhat, B. Shapiro, S. Sheppard, C. Colijn, and M. Murray, “A phylogeny-based sampling strategy and power calculator informs genome-wide associations study design for microbial pathogens,” Genome Med., vol. 6, no. 11, p. 101, 2014.

15. M. Laabei, M. Recker, J. K. Rudkin, M. Aldeljawi, Z. Gulay, T. J. Sloan, P. Williams, J. L. Endres, K. W. Bayles, P. D. Fey, V. K. Yajjala, T. Widhelm, E. Hawkins, K. Lewis, S. Parfett, L. Scowen, S. J. Peacock, M. Holden, D. Wilson, T. D. Read, J. van den Elsen, N. K. Priest, E. J. Feil, L. D. Hurst, E. Josefsson, and R. C. Massey, “Predicting the virulence of MRSA from its genome sequence,” Genome Res., vol. 24, pp. 839–849, May 2014.

16. C. Chewapreecha, P. Marttinen, N. J. Croucher, S. J. Salter, S. R. Harris, A. E. Mather, W. P. Hanage, D. Goldblatt, F. H. Nosten, C. Turner, P. Turner, S. D. Bentley, and J. Parkhill, “Comprehensive identification of single nucleotide polymorphisms associated with beta-lactam resistance within pneumococcal mosaic genes,” PLoS Genet., vol. 10, p. e1004547, Aug. 2014.

17. M. T. Alam, R. A. Petit, 3rd, E. K. Crispell, T. A. Thornton, K. N. Conneely, Y. Jiang, S. W. Satola, and T. D. Read, “Dissecting vancomycin-intermediate resistance in staphylococcus aureus using genome-wide association,” Genome Biol. Evol., vol. 6, pp. 1174–1185, May 2014.

18. M. Wozniak, J. Tiuryn, and L. Wong, “GWAMAR: genome-wide assessment of mutations associated with drug resistance in bacteria,” BMC Genomics, vol. 15 Suppl 10, p. S10, 12 Dec. 2014.

19. K. J. Howell, L. A. Weinert, R. R. Chaudhuri, S.-L. Luan, S. E. Peters, J. Corander, D. Harris, Ø. Angen, V. Aragon, A. Bensaid, S. M. Williamson, J. Parkhill, P. R. Langford, A. N. Rycroft, B. W. Wren, M. T. G. Holden, A. W. Tucker, D. J. Maskell, and BRADP1T Consortium, “The use of genome wide association methods to investigate pathogenicity, population structure and serovar in haemophilus parasuis,” BMC Genomics, vol. 15, p. 1179, 24 Dec. 2014.

20. B. G. Hall, “SNP-associations and phenotype predictions from hundreds of microbial genomes without genome alignments,” PLoS One, vol. 9, p. e90490, 28 Feb. 2014.

21. P. E. Chen and B. J. Shapiro, “The advent of genome-wide association studies for bacteria,” Curr. Opin. Microbiol., vol. 25, pp. 17–24, 25 Mar. 2015.

22. O. Brynildsrud, J. Bohlin, L. Scheffer, and V. Eldholm, “Rapid scoring of genes in microbial pan-genome-wide association studies with scoary,” Genome Biol., vol. 17, p. 238, 25 Nov. 2016.

23. S. G. Earle, C.-H. Wu, J. Charlesworth, N. Stoesser, N. C. Gordon, T. M. Walker, C. C. A. Spencer, Z. Iqbal, D. A. Clifton, K. L. Hopkins, N. Woodford, E. G. Smith, N. Ismail, M. J. Llewelyn, T. E. Peto, D. W. Crook, G. McVean, A. S. Walker, and D. J. Wilson, “Identifying lineage effects when controlling for population structure improves power in bacterial association studies,” Nat Microbiol, vol. 1, p. 16041, 4 Apr. 2016.

24. J. A. Lees, M. Vehkala, N. Välimäki, S. R. Harris, C. Chewapreecha, N. J. Croucher, P. Marttinen, M. R. Davies, A. C. Steer, S. Y. C. Tong, A. Honkela, J. Parkhill, S. D. Bentley, and J. Corander, “Sequence element enrichment analysis to determine the genetic basis of bacterial phenotypes,” Nat. Commun., vol. 7, p. 12797, 16 Sept. 2016.

25. R. A. Power, S. Davaniah, A. Derache, E. Wilkinson, F. Tanser, R. K. Gupta, D. Pillay, and T. de Oliveira, “Genome-Wide association study of HIV whole genome sequences validated using drug resistance,” PLoS One, vol. 11, p. e0163746, 27 Sept. 2016.

26. X. Didelot, D. Lawson, A. Darling, and D. Falush, “Inference of homologous recombination in bacteria using whole-genome sequences,” Genetics, vol. 186, pp. 1435–1449, Dec. 2010.

27. X. Didelot and M. C. J. Maiden, “Impact of recombination on bacterial evolution,” Trends Microbiol., vol. 18, pp. 315–322, July 2010.

28. G. Vernikos, D. Medini, D. R. Riley, and H. Tettelin, “Ten years of pan-genome analyses,” Curr. Opin. Microbiol., vol. 23, pp. 148–154, 2015.

29. M. A. Ansari and X. Didelot, “Bayesian inference of the evolution of a phenotype distribution on a phylogenetic tree,” Genetics, vol. 204, pp. 89–98, Sept. 2016.

30. A. L. Price, N. J. Patterson, R. M. Plenge, M. E. Weinblatt, N. A. Shadick, and D. Reich, “Principal components analysis corrects for stratification in genome-wide association studies,” Nat. Genet., vol. 38, pp. 904–909, Aug. 2006.

31. R. A. Power, J. Parkhill, and T. de Oliveira, “Microbial genome-wide association studies: lessons from human GWAS,” Nat. Rev. Genet., vol. 18, pp. 41–50, Jan. 2017.

32. N. Mantel, “Chi-Square tests with one degree of freedom; extensions of the Mantel-Haenszel procedure,” J. Am. Stat. Assoc., vol. 58, no. 303, pp. 690–700, 1963.

33. K. Pearson, “On lines and planes of closest fit to systems of points in space,” Philosophical Magazine Series 6, vol. 2, no. 11, pp. 559–572, 1901.

34. T. Jombart, S. Devillard, and F. Balloux, “Discriminant analysis of principal components: a new method for the analysis of genetically structured populations,” BMC Genet., vol. 11, p. 94, 15 Oct. 2010.

35. L. A. Weinert, R. R. Chaudhuri, J. Wang, S. E. Peters, J. Corander, T. Jombart, A. Baig, K. J. Howell, M. Vehkala, N. Välimäki, D. Harris, T. T. B. Chieu, N. Van Vinh Chau, J. Campbell, C. Schultsz, J. Parkhill, S. D. Bentley, P. R. Langford, A. N. Rycroft, B. W. Wren, J. Farrar, S. Baker, N. T. Hoa, M. T. G. Holden, A. W. Tucker, D. J. Maskell, and BRaDP1T Consortium, “Genomic signatures of human and animal disease in the zoonotic pathogen streptococcus suis,” Nat. Commun., vol. 6, p. 6740, 31 Mar. 2015.

36. X. Didelot and D. Falush, “Inference of bacterial microevolution using multilocus sequence data,” Genetics, vol. 175, pp. 1251–1266, Mar. 2007.

37. R. Sokal and C. Michener, “A statistical method for evaluating systematic relationships,” University of Kansas Science Bulletin, vol. 38, pp. 1409–1438, 1958.

38. O. Gascuel, “BIONJ: an improved version of the NJ algorithm based on a simple model of sequence data,” Mol. Biol. Evol., vol. 14, pp. 685–695, July 1997.

39. N. Saitou and M. Nei, “The neighbor-joining method: a new method for reconstructing phylogenetic trees,” Mol. Biol. Evol., vol. 4, pp. 406–425, 1 July 1987.

40. A. Criscuolo and O. Gascuel, “Fast NJ-like algorithms to deal with incomplete distance matrices,” BMC Bioinformatics, vol. 9, p. 166, 26 Mar. 2008.

41. J. Felsenstein, “Evolutionary trees from DNA sequences: a maximum likelihood approach,” J. Mol. Evol., vol. 17, no. 6, pp. 368–376, 1981.

42. X. Didelot and D. J. Wilson, “ClonalFrameML: efficient inference of recombination in whole bacterial genomes,” PLoS’ Comput. Biol., vol. 11, p. e1004041, Feb. 2015.

43. N. J. Croucher, A. J. Page, T. R. Connor, A. J. Delaney, J. A. Keane, S. D. Bentley, J. Parkhill, and S. R. Harris, “Rapid phylogenetic analysis of large samples of recombinant bacterial whole genome sequences using gubbins,” Nucleic Acids Res., vol. 43, p. e15, 18 Feb. 2015.

44. W. M. Fitch, “Toward defining the course of evolution: Minimum change for a specific tree topology,” Syst. Biol., vol. 20, pp. 406–416, 1 Dec. 1971.

45. D. L. Swofford and W. P. Maddison, “Reconstructing ancestral character states under wagner parsimony,” Math. Biosci., vol. 87, no. 2, pp. 199–229, 1987.

46. M. Pagel, “Detecting correlated evolution on phylogenies: A general method for the comparative analysis of discrete characters,” Proceedings of the Royal Society of London B: Biological Sciences, vol. 255, pp. 37–45, 22 Jan. 1994.

47. J. Felsenstein, “Maximum-likelihood estimation of evolutionary trees from continuous characters,” Am. J. Hum. Genet., vol. 25, pp. 471–492, Sept. 1973.

48. S. K. Sheppard, X. Didelot, G. Meric, A. Torralbo, K. A. Jolley, D. J. Kelly, S. D. Bentley, M. C. J. Maiden, J. Parkhill, and D. Falush, “Genome-wide association study identifies vitamin B5 biosynthesis as a host specificity factor in campylobacter,” Proc. Natl. Acad. Sci. U. S. A., vol. 110, pp. 11923–11927, 16 July 2013.

49. S. Purcell, B. Neale, K. Todd-Brown, L. Thomas, M. A. R. Ferreira, D. Bender, J. Maller, P. Sklar, P. I. W. de Bakker, M. J. Daly, and P. C. Sham, “PLINK: a tool set for whole-genome association and population-based linkage analyses,” Am. J. Hum. Genet., vol. 81, pp. 559–575, Sept. 2007.

50. N. Patterson, A. L. Price, and D. Reich, “Population structure and eigenanalysis,” PLoS Genet., vol. 2, p. e190, Dec. 2006.

51. N. Mantel and W. Haenszel, “Statistical aspects of the analysis of data from retrospective studies of disease,” J. Natl. Cancer Inst., vol. 22, pp. 719–748, Apr. 1959.

52. C. J. V. Rijsbergen, Information Retrieval. Newton, MA, USA: Butterworth-Heinemann, 2nd ed., 1979.

53. P. B. Frandsen, B. Calcott, C. Mayer, and R. Lanfear, “Automatic selection of partitioning schemes for phylogenetic analyses using iterative k-means clustering of site rates,” BMC Evol. Biol., vol. 15, p. 13, 10 Feb. 2015.

54. C. Tian, P. K. Gregersen, and M. F. Seldin, “Accounting for ancestry: population substructure and genome-wide association studies,” Hum. Mol. Genet., vol. 17, pp. R143–50, 15 Oct. 2008.

55. K. A. Jolley and M. C. J. Maiden, “BIGSdb: Scalable analysis of bacterial genome variation at the population level,” BMC Bioinformatics, vol. 11, no. 1, p. 595, 2010.

56. M. Vos and X. Didelot, “A comparison of homologous recombination rates in bacteria and archaea,” ISME J., vol. 3, pp. 199–208, 2 Oct. 2008.

57. C. Collins and X. Didelot, “Reconstructing the ancestral relationships between bacterial pathogen genomes,” Methods Mol. Biol., vol. 1535, pp. 109–137, 2017.

58. B. A. Oppenheim, “Antibiotic resistance in neisseria meningitidis,” Clin. Infect. Dis., vol. 24 Suppl 1, pp. S98–101, Jan. 1997.

59. L. D. Bowler, Q. Y. Zhang, J. Y. Riou, and B. G. Spratt, “Interspecies recombination between the pena genes of neisseria meningitidis and commensal neisseria species during the emergence of penicillin resistance in n. meningitidis: natural events and laboratory simulation,” J. Bacteriol., vol. 176, pp. 333–337, Jan. 1994.

60. M. C. Maiden, “Horizontal genetic exchange, evolution, and spread of antibiotic resistance in bacteria,” Clin. Infect. Dis., vol. 27 Suppl 1, pp. S12–20, Aug. 1998.

61. B. G. Spratt, Q. Y. Zhang, D. M. Jones, A. Hutchison, J. A. Brannigan, and C. G. Dowson, “Recruitment of a penicillin-binding protein gene from neisseria flavescens during the emergence of penicillin resistance in neisseria meningitidis,” Proc. Natl. Acad. Sci. U. S. A., vol. 86, pp. 8988–8992, Nov. 1989.

62. Q. Y. Zhang, D. M. Jones, J. A. Sáez Nieto, E. Pérez Trallero, and B. G. Spratt, “Genetic diversity of penicillin-binding protein 2 genes of penicillin-resistant strains of neisseria meningitidis revealed by fingerprinting of amplified DNA,” Antimicrob. Agents Chemother., vol. 34, pp. 1523–1528, Aug. 1990.

63. T. Read and R. Massey, “Characterizing the genetic basis of bacterial phenotypes using genome-wide association studies: a new direction for bacteriology,” Genome Med., vol. 6, no. 11, p. 109, 2014.

64. M. Pizza and R. Rappuoli, “Neisseria meningitidis: pathogenesis and immunity,” Curr. Opin. Microbiol., vol. 23, pp. 68–72, Feb. 2015.

65. B. Capecchi, J. Adu-Bobie, F. Di Marcello, L. Ciucchi, V. Masignani, A. Taddei, R. Rappuoli, M. Pizza, and B. Aricò, “Neisseria meningitidis NadA is a new invasin which promotes bacterial adhesion to and penetration into human epithelial cells,” Mol. Microbiol., vol. 55, pp. 687–698, Feb. 2005.

66. M. Comanducci, S. Bambini, B. Brunelli, J. Adu-Bobie, B. Aricò, B. Capecchi, M. M. Giuliani, V. Masignani, L. Santini, S. Savino, D. M. Granoff, D. A. Caugant, M. Pizza, R. Rappuoli, and M. Mora, “NadA, a novel vaccine candidate of neisseria meningitidis,” J. Exp. Med., vol. 195, pp. 1445–1454, 3 June 2002.

67. L. Fagnocchi, E. Pigozzi, V. Scarlato, and I. Delany, “In the NadR regulon, adhesins and diverse meningococcal functions are regulated in response to signals in human saliva,” J. Bacteriol., vol. 194, pp. 460–474, Jan. 2012.

68. S. D. Bentley, G. S. Vernikos, L. A. S. Snyder, C. Churcher, C. Arrowsmith, T. Chillingworth, A. Cronin, P. H. Davis, N. E. Holroyd, K. Jagels, M. Maddison, S. Moule, E. Rabbinowitsch, S. Sharp, L. Unwin, S. Whitehead, M. A. Quail, M. Achtman, B. Barrell, N. J. Saunders, and J. Parkhill, “Meningococcal genetic variation mechanisms viewed through comparative analysis of serogroup C strain FAM18,” PLoS Genet., vol. 3, p. e23, 16 Feb. 2007.

69. O. B. Harrison, N. J. Evans, J. M. Blair, H. S. Grimes, C. R. Tinsley, X. Nassif, P. Kriz, R. Ure, S. J. Gray, J. P. Derrick, M. C. J. Maiden, and I. M. Feavers, “Epidemiological evidence for the role of the hemoglobin receptor, hmbr, in meningococcal virulence,” J. Infect. Dis., vol. 200, pp. 94–98, 1 July 2009.

70. I. Stojiljkovic, V. Hwa, L. de Saint Martin, P. O’Gaora, X. Nassif, F. Heffron, and M. So, “The neisseria meningitidis haemoglobin receptor: its role in iron utilization and virulence,” Mol. Microbiol., vol. 15, pp. 531–541, Feb. 1995.

71. I. Stojiljkovic, J. Larson, V. Hwa, S. Anic, and M. So, “HmbR outer membrane receptors of pathogenic neisseria spp.: iron-regulated, hemoglobin-binding proteins with a high level of primary structure conservation,” J. Bacteriol., vol. 178, pp. 4670–4678, Aug. 1996.

72. I. Chen and E. C. Gotschlich, “ComE, a competence protein from neisseria gonorrhoeae with DNA-binding activity,” J. Bacteriol., vol. 183, pp. 3160–3168, May 2001.

73. L. A. S. Snyder, J. A. Cole, and M. J. Pallen, “Comparative analysis of two neisseria gonorrhoeae genome sequences reveals evidence of mobilization of correia repeat enclosed elements and their role in regulation,” BMC Genomics, vol. 10, p. 70, 9 Feb. 2009.

74. D. J. Hill, N. J. Griffiths, E. Borodina, and M. Virji, “Cellular and molecular biology of neisseria meningitidis colonization and invasive disease,” Clin. Sci., vol. 118, pp. 547–564, 9 Feb. 2010.

75. E. Capel, A. L. Zomer, T. Nussbaumer, C. Bole, B. Izac, E. Frapy, J. Meyer, H. Bouzinba-Ségard, E. Bille, A. Jamet, A. Cavau, F. Letourneur, S. Bourdoulous, T. Rattei, X. Nassif, and M. Coureuil, “Comprehensive identification of meningococcal genes and small noncoding RNAs required for host cell colonization,” MBio, vol. 7, 7 Sept. 2016.

76. R. Urwin, J. E. Russell, E. A. L. Thompson, E. C. Holmes, I. M. Feavers, and M. C. J. Maiden, “Distribution of surface protein variants among hyperinvasive meningococci: implications for vaccine design,” Infect. Immun., vol. 72, pp. 5955–5962, Oct. 2004.

77. J. E. Russell, K. A. Jolley, I. M. Feavers, M. C. J. Maiden, and J. Suker, “PorA variable regions of neisseria meningitidis,” Emerg. Infect. Dis., vol. 10, pp. 674–678, Apr. 2004.

78. J. P. Derrick, R. Urwin, J. Suker, I. M. Feavers, and M. C. Maiden, “Structural and evolutionary inference from molecular variation in neisseria porins,” Infect. Immun., vol. 67, pp. 2406–2413, May 1999.

79. J. Suker, I. M. Feavers, M. Achtman, G. Morelli, J. F. Wang, and M. C. Maiden, “The pora gene in serogroup a meningococci: evolutionary stability and mechanism of genetic variation,” Mol. Microbiol., vol. 12, pp. 253–265, Apr. 1994.

80. S. A. Tunio, N. J. Oldfield, D. A. A. Ala’Aldeen, K. G. Wooldridge, and D. P. J. Turner, “The role of glyceraldehyde 3-phosphate dehydrogenase (GapA-1) in neisseria meningitidis adherence to human cells,” BMC Microbiol., vol. 10, p. 280, 9 Nov. 2010.

